# Identifying memory gene expression from single sample scRNA-seq data using power law signatures

**DOI:** 10.1101/2025.01.15.633183

**Authors:** Suvranil Ghosh, Shaon Chakrabarti, Archishman Raju

**Affiliations:** Simons Centre for the Study of Living Machines, National Centre for Biological Sciences, Tata Institute of Fundamental Research, Bangalore 560065, India

## Abstract

Genes with expression levels that fluctuate on time scales longer than cell division times are associated with cancer drug tolerance. However, current methods for identifying such ‘memory’ genes rely on variants of the Luria-Delbrück experiment and require either multiple replicates or lineage information, limiting their use to model systems or *in-vitro* settings. We develop a new conceptual approach using recent results in Random Matrix Theory to demonstrate that the existence of memory genes results in a power-law signature in the cell covariance matrix eigenspectrum. Utilizing this theoretical framework, we develop Power-Seek, an algorithm to discover memory genes from a single time point scRNA-seq dataset. Without using prior information on lineages or cell-cycle times, Power-Seek correctly identifies memory genes in a melanoma cell line. Our results open up the possibility of identifying expression states driving drug tolerance in real-world scenarios, as we demonstrate using data from a human breast cancer tissue sample.

## 2 Introduction

Fluctuations in the levels of various cellular components – RNA, proteins, histone, and DNA methylation – result in the existence of different ‘cell states’ at any given point in time, leading to a variety of dynamical trajectories being followed by different groups of cells. The heterogeneity in cell states within genetically identical cells can also have a major effect on their response to perturbations. Examples where such non-genetic heterogeneity in cell states plays a role include developmental outcomes [1, 2, 3], the response of bacteria to stress [4, 5], proliferation-quiescence decisions at mitosis [6, 7], and cancer cell survival on drug treatment [8, 9, 10]. Discovering these non-genetic states and connecting them to eventual fate remains a challenging problem across various fields of biology [11].

Recent work has brought attention to the inheritability of these cell states, which play a role in cell fate. Cells may drift into rare states as a result of the fluctuation of a few slowly decaying ‘memory genes’ [12, 13]. These memory genes have relaxation times that are long compared to cell cycle times, and hence their expression states are inherited across cell divisions [14, 15]. Interestingly, these memory genes have recently been associated with eventual drug tolerance in cancer [12, 16]. For instance, *EGFR* and *AXL*, well-known genes associated with drug tolerance and cell state transitions, were found to have characteristics of memory genes and primed melanoma cells towards vemurafenib resistance [12, 16]. Typically, memory genes can have coordinated expression giving rise to a memory state [17, 16]. Often, however, the identity of the memory states are unknown even though their association with fate outcomes is clear from correlations between lineage-related cells [8, 18, 19, 20]. For example, post-cisplatin p53 dynamics driving colon-cancer cell death versus survival [9] is strongly correlated in sister cells [21], suggesting upstream control via memory states. However, identifying these memory states priming post-drug p53 responses has been challenging. Discovering these states and the underlying memory genes has therefore emerged as an exciting new direction for understanding and steering cell fate outcomes.

Currently, most methods to estimate whether a gene is a memory gene or not rely on a Luria-Delbrück like method. In lineage related cells, if a rare fluctuation in a gene drives a cell to a particular cell state early in the lineage, all cells in the lineage will have strongly correlated expression in that gene. Variants of this basic idea can be used to identify memory genes [12, 22, 23, 24] and epigenetic states more broadly [25]. In general, single time point measurements can be combined with lineage information to infer the dynamics of cell states [26, 27, 23]. However, except in model systems where genetic engineering is feasible or *in vitro* settings, such lineage information is unlikely to be available.

Here, we introduce a novel theoretical framework to estimate memory genes from single time point measurements of lineage related cells without explicit lineage information. Our framework is based in Random Matrix Theory, which has recently been applied to analyze scRNA-seq data [28, 29, 30]. We build on recent results that use Random Matrix Theory to estimate signatures of lineage correlation in protein sequences [31, 32]. We extend these results to models of continuous gene expression with variable decay rates and develop an algorithm, Power-Seek, to predict the presence of memory genes from single time point measurements. Our framework relies on utilizing the signatures of lineage correlations without direct access to lineage information. The algorithm we develop is justified by our theoretical framework, has essentially no free parameters, and is easy to implement. We first apply Power-Seek to a single time point scRNA-seq dataset from the melanoma cell line WM989 and demonstrate how it is able to robustly identify memory genes previously identified using many independent samples and bulk RNA-sequencing methods. We then apply Power-Seek to identify memory genes from a patient breast cancer tissue dataset. Our results open up the exciting possibility of discovering inheritable gene expression states in clinical settings, not limited by sample numbers or the availability of genetically encoded barcodes.

## 3 Results

### 3.1 Lineages carry dynamic information which can be recovered from a static snapshot

We assume that gene expression, as measured by the mRNA concentration, fluctuates about a steady state. Fig. 1A shows the dynamics of gene expression of two genes if the system is perturbed away from its steady state at some time *t*. Such a perturbation may be externally induced or simply be the result of a large stochastic fluctuation. The expression of each gene will relax back to its steady state value with some typical time scale. For a memory gene, the expression level decays slower than the underlying average cell cycle time *τ_c_* (green line in Fig. 1A) while a ‘non-memory’ gene is one whose expression level decays faster than *τ_c_* (magenta line in Fig. 1A). The degree of inheritance of a memory gene can then be quantified as the half-life of the relaxation dynamics measured in units of cellular generations *m* (Fig. 1B). Non-memory genes have *m <* 1 whereas memory genes have larger values of *m*, and the distribution of half-lives is typically skewed towards non-memory genes [33] (Fig. 1B).

**Figure 1:**
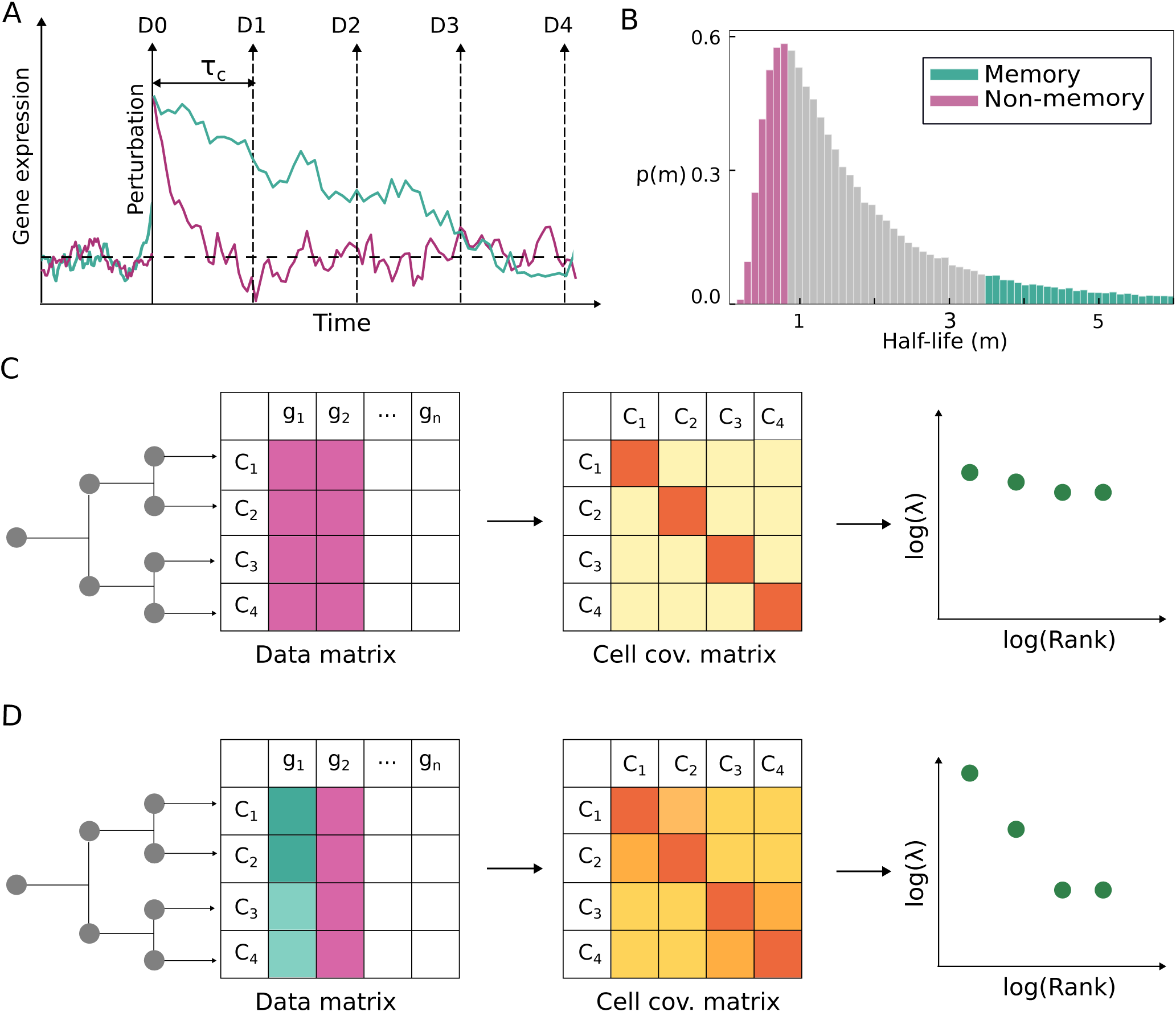
Schematic of the effect of lineage correlation on gene expression dynamics. **A:** Representative schematic of two genes whose expression levels fluctuate around the same steady state (horizontal dashed line). If the genes are perturbed away from the steady state as a result of external changes or a stochastic fluctuation, their expression levels decay at different rates. The vertical dashed lines show cell division events. A memory gene (green) decays slower than the average cell cycle time, while a non-memory gene (magenta) decays to the steady state within one generation. For clarity, it is assumed here that only one daughter cell is tracked at each division, which perfectly inherits the gene expression of the mother. Gene expression is measured as the concentration of mRNA levels. **B:** Distribution of half-lives of gene expression states as measured in units of cellular generations. **C:** A representative lineage with 4 cells produces a data matrix with gene expression values for every cell. This data matrix can be used to construct a cell covariance matrix which will have a flat distribution of eigenvalues if only non-memory genes are present. **D:** If some of the genes are memory genes, the eigenspectrum of the cell covariance matrix widens which is a signature of lineage correlations.

The presence of memory genes implies that cells undergoing division will have correlated gene expression in their lineage. Consider one such lineage as shown in Fig. 1C, where gene expression is measured after a couple of cell divisions. We can define a data matrix ***X*** with cells as rows and gene expression values, centralized about their mean and normalized by the standard deviation across cells, as columns. The data matrix can be multiplied by its transpose and divided by the number of genes to generate a cell covariance matrix, which quantifies the total expression correlation arising from all genes between each pair of single cells. If only non-memory genes are measured, gene expression in sisters, cousins, etc., will not be correlated. The cell covariance matrix will have ones on the diagonal and noise in the off-diagonal elements, resulting in a flat eigenvalue spectrum (Fig. 1C).

Note that because of our normalization, there is no separation in eigenvalues because of differences in variability of genes. However, cells will be correlated in the presence of memory genes, with the extent of correlations depending on the half-life of the memory genes and lineage distance between cells (Fig. 1D). These correlations will typically lead to wider separations in eigenvalues. Crucially, the eigenvalues of the cell covariance matrix are independent of the row ordering in the data matrix. As a result, even if lineage relationships between the cells are unknown, the wide separations of eigenvalues provide a signature of correlations arising from memory genes. A single snapshot can, therefore, in principle, be used to infer the presence of inheritable memory-gene expression states. We will formalize these ideas in the next section.

### 3.2 Inheritable gene expression dynamics generates power laws in the cell covariance matrix eigenspectrum

We now quantify the intuitive results of the previous section with a mathematical model of gene expression dynamics, to demonstrate how the cell covariance matrix eigenspectrum differs in the presence and absence of memory genes. We assume that each gene has an associated relaxation time scale and stochastic fluctuations about a steady state. The precise form of the distribution of relaxation time scales is not crucial for our results (Fig. **SI 1**), and we model it with a Gamma distribution (Methods, Eq. 2). We model gene expression dynamics with an Ornstein-Uhlenbeck process (Methods, Eq. 1). The lineage is assumed to begin with one cell and at every division, each daughter inherits the expression level of the mother cell. Cell division times are taken to be constant so that after *b* generations, there are 2*^b^* cells. The data matrix ***X*** is constructed from these 2*^b^* cells, where the expression levels of each gene are centralized around the mean and normalized by the standard deviation across cells.

The data matrix can be used to construct both a cell covariance and a gene covariance matrix which will have essentially the same eigenvalues. Naively, one might expect that eigenvectors of the gene covariance matrix have information about the memory genes. In the absence of cell-cell correlations, these eigenvectors may indeed have this information [34], but this information is no longer present in the presence of lineage correlations, as we show in the SI Section **S2**.

The cell covariance matrix for our model depends only on a dimensionless parameter given by the ratio of the cell cycle time to the mean relaxation time, which we denote by *τ*. Large values of *τ* indicate that most genes are non-memory genes, whereas small *τ* implies the presence of memory genes in the system.

To compute the eigenspectrum of the cell covariance matrix, a distinction needs to be made between the ‘true’ and ‘sample’ covariance matrices [31]. The true cell covariance matrix is defined as the expected cell covariance for an ensemble. Hence, averaging over many replicates gives the true cell covariance matrix shown in Fig. 2A. The block structure of the true cell covariance matrix reflects the lineage relationships; every cell has one sister, two first cousins etc. similar to Fig 1D. This block structure allows for an analytical calculation of the true covariance matrix as we outline in Methods and SI Section **S3**. In practice, however, gene expression is measured for a single sample, giving rise to the sample covariance matrix shown in Fig. 2B. Further, lineage relationships are typically unknown; hence, the sample covariance matrix looks noisy and without structure.

**Figure 2:**
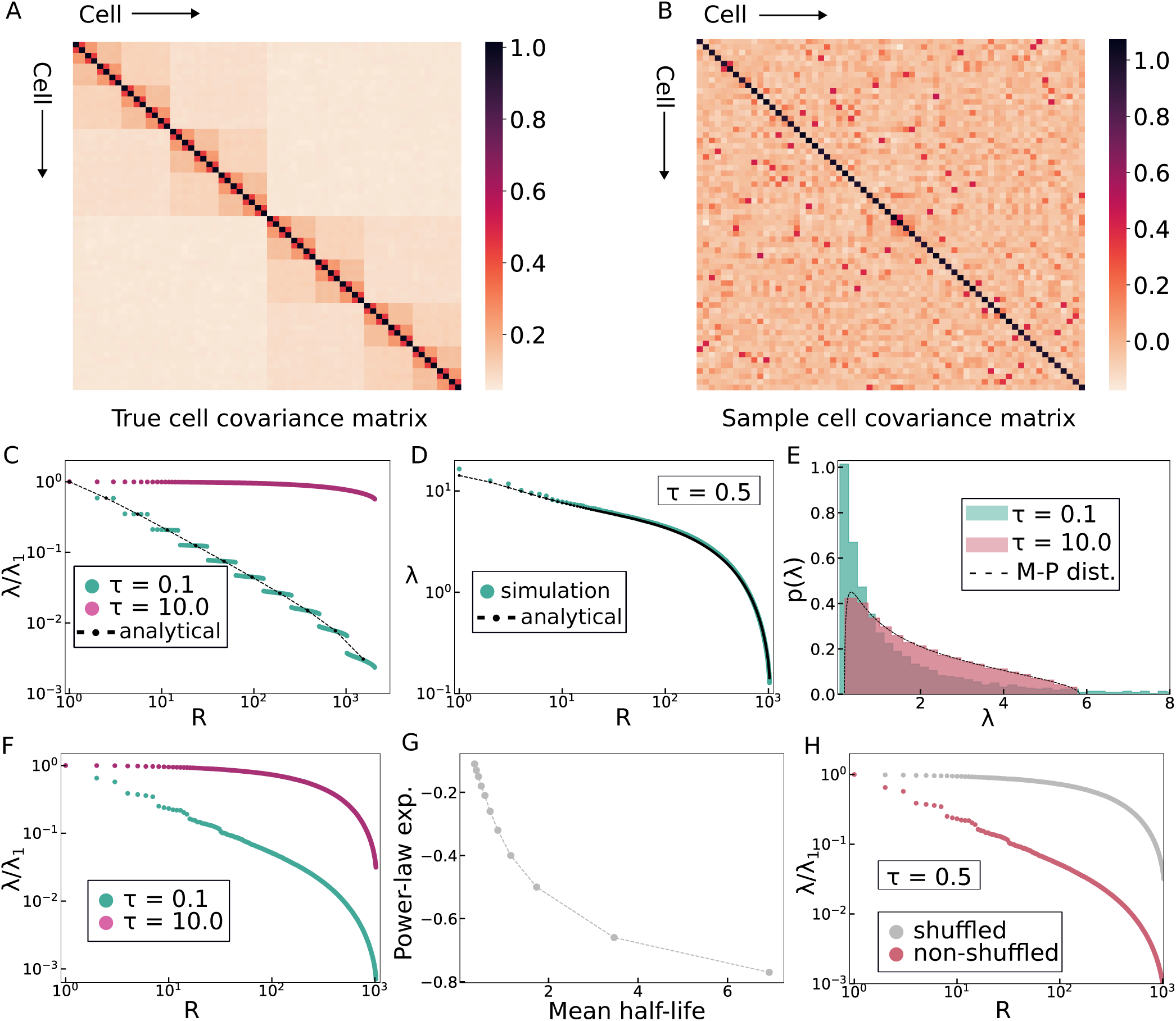
Gene expression dynamics with lineage correlation leads to power law in the eigenspectra of the cell covariance matrix. **A**: True cell covariance matrix. Here, *n_c_* = 64, *n_g_* = 1024, *τ* = 0.5, replicates = 100. Lineage correlation decreases with the lineage distance between two cells giving rise to the visible block structure in the matrix. **B**: Sample covariance matrix with a random ordering of cells. The same parameters are used as in Fig. 2A with a single replicate. **C**: Eigenvalue vs. Rank curve for true covariance matrix for different mean relaxation times. The eigenvalues are degenerate and black dots show our analytical solution (Methods Eq. 4) with the dashed line drawn as a guide to the eye. **D**: Eigenvalue vs. Rank curve for a single sample from simulation and analytics. The black dots show our analytical solution (SI Section S4) with the dashed line drawn as a guide to the eye. **E**: Histogram of (non-zero) eigenvalues from a single snapshot for different mean half-lives. The Marchenko-Pastur (M-P) distribution (Methods Eq. 5) is plotted as a dashed line. **F**: Eigenvalue vs. Rank curve from a single snapshot for different mean half-lives. **G**: A plot of the power law exponent versus the mean half-lives (in units of cellular generations). The power law exponent is negative and has a maximum magnitude of one. Small exponents reflect less memory. **H**: Eigenvalues vs Rank curve for the shuffled and non-shuffled data matrix.

The eigenspectra of the true covariance matrix for two different values of *τ* are shown in Fig. 2C. The plot of the eigenvalues with rank for low values of *τ* shows degenerate eigenvalues with a power law structure, which is a signature of lineage correlations. This power law formalizes the notion of separation of eigenvalues mentioned in the previous section. On the other hand, high values of *τ* show a relatively flat spectrum, and the distribution of eigenvalues is predicted to be the Marchenko-Pastur distribution [35].

We extend the analytical calculations to predict the distribution of eigenvalues for a single sample using a combination of results from Random Matrix Theory and order statistics (see Methods and SI Section **S4**). A comparison of our analytical results with numerical simulations is shown in Fig. 2D. The eigenvalues do not depend on the ordering of rows in the data matrix and hence Fig. 2D has information about the lineage correlations independent of our knowledge of lineage relationships. In the limit where lineage relationships are no longer present, given by large *τ* in our model, we get a Marchenko-Pastur distribution. The comparison of our analytical and numerical results is shown for large and small *τ* in Fig. 2E. Our results generalize and extend the results of [31] to models of continuous gene expression. In a single sample, unlike the eigenspectrum of the true covariance matrix, only a fraction of eigenvalues follow a power law followed by a non power law regime (Fig. 2F). The eigenvalues within the power law regime are the ones that carry information on lineage correlations.

Additionally, we find that the power law exponent is informative about the average number of generations for which gene expression values remain correlated where the average is computed over all genes. Analytical calculations simplify for small *τ* (SI Section **S**3.3), and we show numerical results in Fig. 2G. Finally, we verify that the power laws are a signature of correlations by shuffling gene expression values between randomly selected cells (details in Methods), which gives back the Marchenko-Pastur distribution (Fig 2H).

### 3.3 Power-Seek: An algorithm to identify memory genes from single sample scRNA-seq data

We now develop an algorithm to detect memory genes from an scRNA-seq dataset building on the theoretical results of the last section. As we demonstrated, the power law in the eigenvalue spectrum of the cell covariance matrix results from the lineage correlations driven by memory genes. Consequently, if a gene is removed from the data matrix we expect the direction of change in the eigenvalues spectrum to be different depending on whether the gene is a memory or non-memory gene. Removing a non-memory gene has a small effect on the spectrum, whereas removing a memory gene has a larger effect, reducing the separation between eigenvalues as shown in Fig 3A. We utilize this idea to remove a single gene at a time from the data matrix and classify the removed gene as memory or not, based on the direction of change of the lineage-related eigenvalues (Fig 3A).

**Figure 3:**
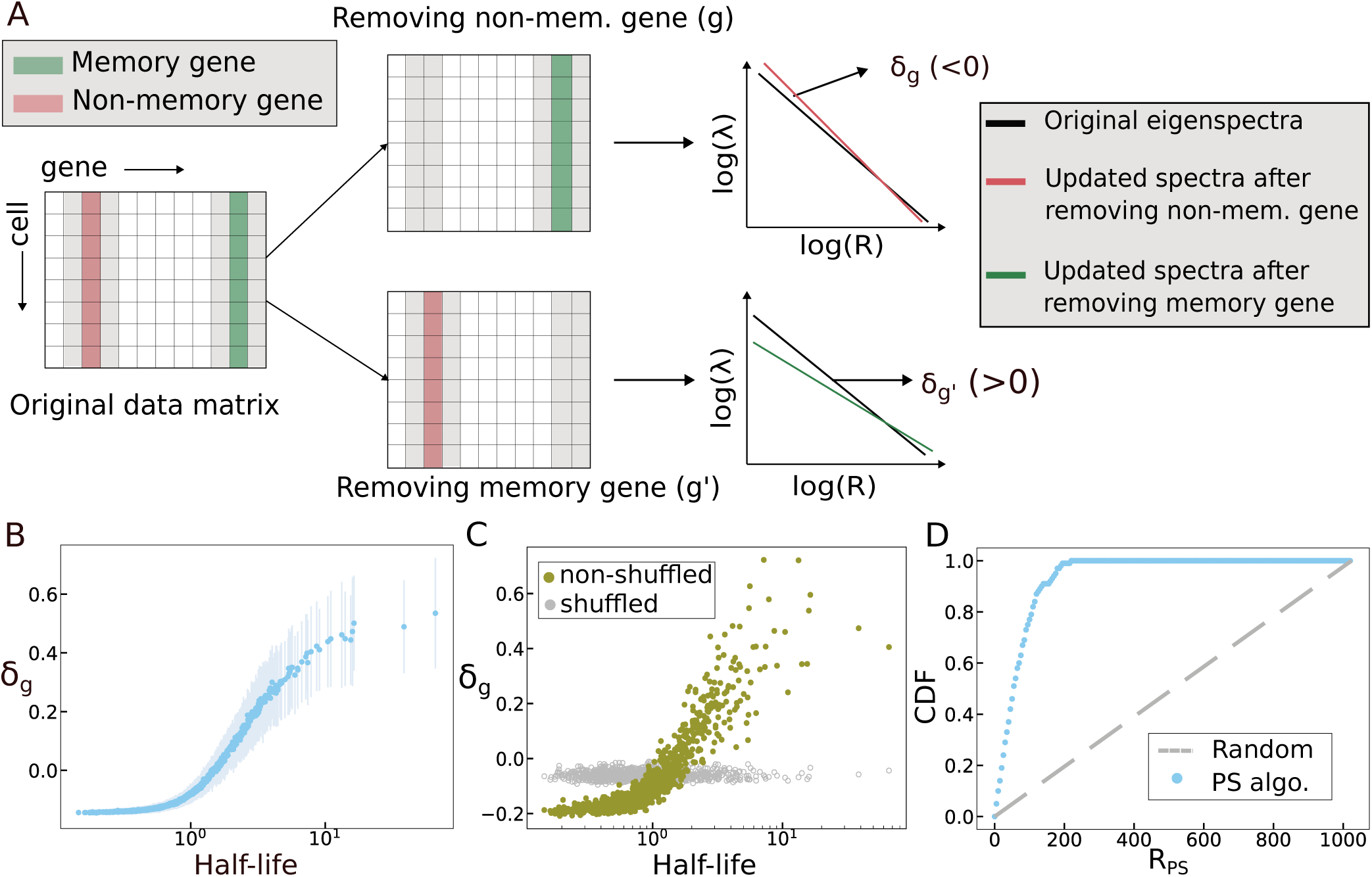
Power-Seek algorithm estimates the ordering of intrinsic relaxation timescales for different genes from the data matrix. **A**: Flowchart of Power-Seek algorithm. A single gene is removed from the data matrix. If a non-memory gene is removed, it leads to a small negative change in the eigenvalue spectrum as measured by *δ_g_* (defined in main text). A memory gene leads to a larger positive change as the distribution becomes more like the Marchenko-Pastur distribution. **B**: A plot of *δ_g_* versus the half-life as measured in units of cell cycle time shows strongly monotonic behavior. The gray lines indicate the variability from sample to sample, and the blue dots are the mean across samples. Parameters: *n_c_* = 2048, *n_g_* = 1024, *τ* = 0.5, number of samples = 40. **C:** For the same parameter set as (B), the quantity *δ_g_* plotted against the half-life for a single sample. Grey dots show the behavior of *δ_g_* when the Power-Seek algorithm is applied to a shuffled simulation data matrix. **D**: Proportion of top 100 ‘true memory genes’ (as measured by our relaxation time scales) as a function of rank predicted by the Power-Seek algorithm. The gray dashed line denotes the expected number of genes by chance.

To this end, we define a quantity *δ_g_*, which is the difference of the eigenvalues between the covariance matrices resulting from the new data matrix (obtained by removing a single gene *g*) and the original data matrix. Therefore *δ_g_* for a specific gene *g* is given by 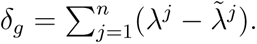 and 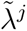 are the *j^th^* rank eigenvalues of the original and new cell covariance matrices respectively. The threshold number of eigenvalues *n* is estimated by the number of eigenvalues that are in the power law regime (details in Methods). In Fig. 3B, we show that *δ_g_*is very strongly correlated with the half-life of that particular gene. Furthermore, even though there is sample to sample variability in *δ_g_* for any particular gene *g* (Fig. 3B), memory genes with larger half-life will typically have *δ_g_ >* 0 while nonmemory genes will have *δ_g_ <* 0. We examine the dependence of *δ_g_* on the number of cells and genes in SI Section **S**6.

The quantity *δ_g_* does not give any information if the data matrix is shuffled in the same way as before, which we verify in Fig. 3C. While we can not directly calculate *δ_g_* analytically, we can approximate it, and our analytical results are qualitatively very similar to our numerical results, as shown in Fig. **SI** 3. Our analytical calculations show why, for non-memory genes, *δ_g_* goes to a small negative number rather than zero (SI Section **S**5).

It is possible to visualize the efficacy of the Power-Seek algorithm in another way. If we select the top 100 memory genes as defined by the relaxation time scales, we can plot the cumulative distribution function of the probability of finding these memory genes as a function of Power Seek rank as done in Fig 3D. The sharply rising behavior of this function, as opposed to the linear behavior expected if memory genes were being found randomly, demonstrates the algorithm’s efficacy in identifying memory genes.

### 3.4 Power-Seek predicts memory genes in melanoma cells from a single sample scRNA-seq dataset

Having demonstrated that Power-Seek can correctly identify memory genes in simulated datasets, we next explore if Power-Seek can recover previously identified memory genes from an scRNA-seq dataset. We test our results using data in [12] and [36]. The first of these performed bulk RNA sequencing on barcoded cells in the melanoma cell line WM989 to identify memory genes using the Luria-Delbrück test. Ref. [36] performed scRNA-seq on the same cell line. We identify memory genes from the scRNA-seq data and compare them to the memory genes found in [12].

We obtain a cell covariance matrix after preprocessing the data (Methods) from each of the two replicates available. The theoretical framework outlined in the previous sections predicts that the eigenvalues of the cell covariance matrix will have a power law form. These eigenvalues are shown in Fig. 4A and have a power law structure as predicted, for both replicates. We use only the first replicate for the rest of our analysis with the analysis for the second replicate shown in the SI Section **S**7. In Fig. 4B, we verify that if we shuffle the data, we do not get any power law structure in the eigenvalues. Therefore, we conclude that lineage correlations in the data will allow us to identify the underlying memory genes using the algorithm.

**Figure 4:**
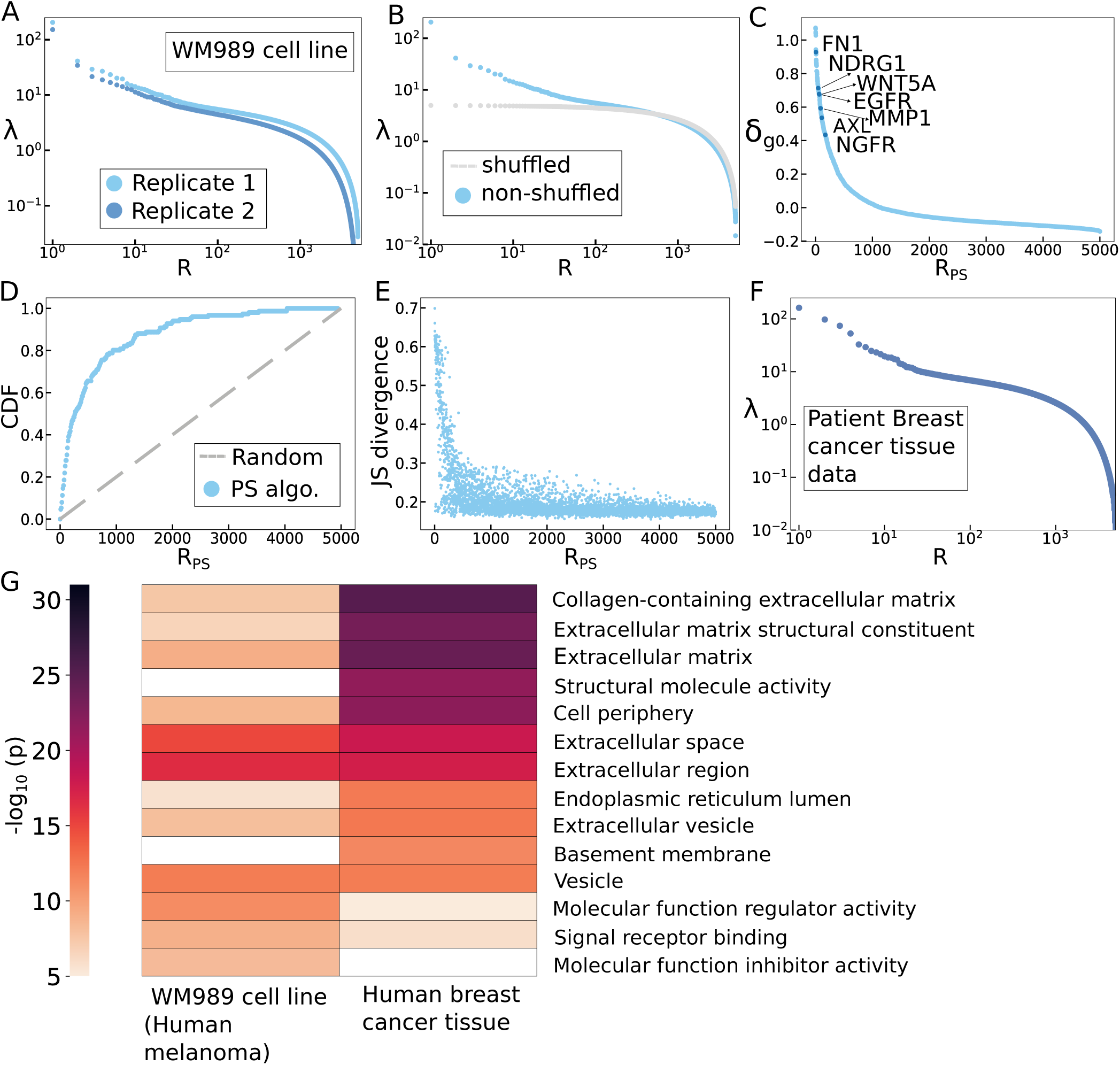
Power-Seek algorithm predicts memory genes using scRNA data from WM989 melanoma cell line. **A:** The eigenvalues of the cell covariance matrix for scRNA-seq data from a melanoma cell line show a power law structure. The eigenvalues are plotted for two replicates. In replicate 1, *n_c_* = 7590, *n_g_* = 3000 and in replicate 2, *n_c_* = 4941, *n_g_* = 5000. **B:** A comparison of the original eigenvalue spectra of the cell covariance matrix with the spectra of the cell covariance matrix obtained by shuffling the data matrix. The spectra of the shuffled matrix show no power laws. **C:** A plot of the difference metric *δ_g_* with Power-Seek Rank *R_P S_*. Higher *δ_g_* denotes higher memory. i.e., lower Power-Seek rank *R_PS_*. The positions of memory genes in [12] are marked. **D:** Overlap between memory genes found out by Power-Seek algorithm and MemorySeq. The gray dashed line denotes the expected number of genes by chance. **E:** A plot of the Jensen-Shannon divergence between the distribution of gene expression in a cluster with plausible lineage relationship and the distribution of gene expression across all cells. The cluster was formed using the 1000 cells with the highest gene expression cells of a memory gene as identified by Power-Seek. **F:** The eigenvalues of the cell covariance matrix for scRNA-seq data from a patient breast cancer tissue data. Here, *n_c_* = 7967, *n_g_* = 5000. **G:** The heatmap shows the functional categories that are most prevalent in the two data sets. *p* denotes the probability of finding genes associated with those functional categories by chance and darker colours indicate overrepresented categories. In both data sets, genes associated with the extracellular matrix are in the top overrepresented categories.

We analyze the cell covariance matrix using our Power-Seek algorithm and plot sorted values of *δ_g_* in Fig. 4C. We find several memory genes, as identified in [12], which have high values of *δ_g_*. Out of the 227 memory genes identified in [12], 151 are present in our data matrix after all pre-processing steps. We find 99 of these 151 memory genes (about 65%) in the top 500 Power-Seek rank genes. Ref. [12] had also tested the inheritance of a few of their predicted memory genes using time-lapse microscopy and RNA FISH. Importantly, all these validated genes appear in the top 200 of our predicted memory genes from the Power-Seek algorithm.

To further demonstrate that Power-Seek is not simply a random classifier of memory versus nonmemory genes, we plot the proportion of memory genes in [12] identified by Power-Seek as a function of Power-Seek rank. If memory genes were being identified at random, we would expect a straight line (Fig. 4D). On the contrary, we get a sharply rising curve, which suggests that Power-Seek is able to correctly find a large proportion of memory genes.

We can independently test that the genes that we identify are memory genes using the following procedure. First, we select a memory gene as calculated by our algorithm and look at a cluster of cells with high expression of that memory gene. This cluster of cells is, therefore, likely to be related by lineage. For all genes, we then measure the Jensen-Shannon divergence of the distribution of gene expression in this cluster and the gene expression across all cells. This is a measure of how different the gene expression is in cells that likely have lineage correlations as compared to the gene expression in all cells. We see a strong correlation between Jensen-Shannon divergence and Power Seek rank (Fig. 4E), suggesting that the top-ranked genes are indeed those that carry memory.

### 3.5 Memory genes in a patient breast cancer tissue sample are enriched for extracellular matrix and EMT functions

Having verified that Power-Seek can reproducibly identify known memory genes in two independent scRNA-seq replicates from the WM989 melanoma cell line, we next explore its applicability in a patient tissue sample. We anticipate Power-Seek to be particularly useful in clinical settings, where obtaining multiple samples or barcoding via genetic engineering is not feasible and hence approaches using the Luria-Delbrück fluctuation test would not be possible. Here, we apply Power-Seek to an scRNA-seq data from a formalin-fixed, paraffin-embedded (FFPE) patient breast cancer tissue block (TNM stage T2N1M0) [37].

After preprocessing the scRNA-seq data (see Methods), we calculate the cell covariance matrix, whose eigenvalues reveal a power law structure as shown in Fig. 4F. This suggests the existence of memory genes in the breast cancer tissue data, and the possibility of detecting them using Power-Seek. We then apply Power-Seek as before, and identify the top ranked memory genes. To compare memory genes from the WM989 cell line and patient breast cancer tissue, we perform over-representation analysis using Gene Ontology Cellular Components and Molecular Functions datasets [38] (see Methods). Fig. 4G shows the combined list of the top 10 functional categories for WM989 cell lines and patient tissue. Notably, ‘extracellular matrix’ (ECM) genes are prominent in most overrepresented functional categories for both, consistent with independent studies linking these genes to lineage-related cells across samples [39]. Previous work has suggested that ECM-related genes might carry micro-environmental information and be used as biomarkers of stable cell states [40].

Interestingly, we also find that among the top 200 memory genes in the breast cancer tissue, 39 genes belong to Epithelial-Mesenchymal Transition (EMT)-associated hallmark genes as per the Molecular Signatures Database [41], while only 78 are present among the top 5000 highly variable genes. Similarly, in the WM989 cell line (replicate 1), we find 32 EMT-associated hallmark genes in the list of top 200 memory genes among a total of 77 EMT-associated genes among the top 5000 highly variable genes. Among the top 200 memory genes in both WM989 melanoma and breast cancer tissue, we find many invasiveness genes like *FLT1*, *EGFR*, *AXL*, *FN1*, *VEGFC*, *PDGFC*, *NGFR*, *ZEB1*, etc [42, 43]. These results suggest that EMT gene expression states, often associated with drug resistance and metastasis, are likely inheritable over cellular generations across cancer types.

## 4 Discussion

In this work we develop a novel theoretical approach to demonstrate that memory genes – a class of genes whose expression levels are inheritable across a few cellular generations, can be identified from a single sample, single time point scRNA-seq dataset without requiring barcodes or any prior knowledge of the underlying cell cycle times. While this result is rather counter-intuitive, we show that it is feasible because of a subtle underlying feature of memory genes – they generate expression correlations in lineage-related cells, which in turn results in a power law in the eigenspectrum of the experimentally measured cell covariance matrix. Importantly, the power law exponent is related to the memory timescale – genes whose expression states are more inheritable give rise to larger exponents. Therefore, even if the underlying lineage structure among single cells is unknown, the presence of a power law and its dependence on inheritable gene expression can be utilized to discover the underlying memory genes. We rigorously demonstrate these ideas by extending a previous result in the context of protein evolution [31]. We model continuous gene expression dynamics, and provide new theoretical results for the power law behavior and its relationship with memory timescales. We then leverage these conceptual ideas to develop Power-Seek, a computational algorithm that discovers memory genes from experimental scRNA-seq datasets.

Our results open up the possibility of using a single sample to identify genes related to drug tolerance in cancer. To demonstrate the ability of Power-Seek to identify previously discovered and validated inheritable genes, we analyze an scRNA-seq dataset from the WM989 melanoma cell line. We recover 65% of the memory genes that were originally discovered from 43 samples using a Luria-Delbrück fluctuation test. Indeed, genes like *FN1, EGFR, NGFR* and *AXL*, which were validated using smFISH and considered strong memory genes, all showed up as top ranked Power-Seek genes. To then demonstrate Power-Seek’s ability to discover memory genes where other Luria-Delbrück based approaches are unlikely to work due to unavailability of multiple samples or barcodes, we analyze a single scRNA-seq dataset from a human breast cancer tissue. Interestingly, like in the melanoma dataset, we find that Power-Seek discovers genes that are enriched for extracellular matrix (ECM), cell adhesion and EMT-related functions. This is consistent with independent studies that discovered a role for the ECM in lineage related barcoded cells ([39, 16]). Our results are also consistent with a plethora of previous results linking EMT to breast cancer both *in vitro* and *in vivo* [43, 44, 45]. EMT-associated genes play important roles in increasing cancer cell motility, double-strand DNA repair and escaping oncogene induced senescence in many epithelial (like breast and colon cancer) and non-epithelial (like melanoma) tumor types [45, 25, 42].

Discovery of non-genetic cell states that drive cell fate outcomes is a common challenge across many fields of biology. Recent developments in multiplexed single-cell ‘omics’ technologies provide only static snapshots from which the underlying dynamics of cell states need to be inferred. This is a challenging task requiring the development of careful theoretical methods, and the kinds of dynamics that can be inferred from a single time point dataset remain an open question. Our results on identifying memory genes add to a growing field of identifying dynamics from high-throughput snapshot measurements. However, they are distinct from the more extensively explored question of predicting the future state of a cell using ‘pseudo-time’ [46, 47] or ‘RNA velocity’ methods [48]. The major distinguishing factor is that for memory genes, the expression dynamics are defined with respect to the underlying cell cycle time, and our theoretical approach relies on the gene expression decay timescale being slower than the average cell cycle time. The fundamental relationship to the cell cycle also raises the intriguing possibility that memory genes may be context-dependent – for example, even for the same cell type, spatial constraints might slow down the cell cycle, leading to an otherwise memory gene converting to a non-memory gene. With the rapid development of spatial transcriptomics, Power-Seek could prove powerful in detecting such context dependence, for example, between the core and periphery of a tumor [49].

Power-Seek has no free parameters aside from a threshold on the number of eigenvalues and is extremely simple to implement. Nevertheless, our theoretical framework is built on certain underlying assumptions and associated limitations. Primary among them is that we do not model multiple states and transitions between them but assume a single stable state with exponential relaxation after a fluctuation. While this assumption may not be the most accurate description of non-genetic cell state dynamics, this is largely how memory gene dynamics have been envisaged in earlier papers and remains an area for future work. The good match between our method and Memory-Seq [33] also suggests that this assumption may not be severely limiting.

It is possible that the power laws observed in the data [36] are due to other underlying factors that cause cell correlations, like spatial signaling. However, obtaining power laws from such a mechanism will typically require fine-tuning, whereas lineage correlations are a robust and natural way to obtain power law correlations. While we can not rule out other mechanisms, our framework relies on a very specific signature of lineage correlations. The breast cancer tissue sample may have multiple cell types and differentially expressed genes may show up as memory genes. Further theoretical and experimental work is required to better understand the role of memory genes in such samples.

While we are able to obtain an ordering of the genes according to their relaxation time scales, we are not able to calculate the number of generations that the memory of any particular gene will last. This means there is a degree of arbitrariness in defining the precise boundary between memory and non-memory genes. In future work, we plan to examine how much information single snapshots give about the relaxation time scales of individual genes.

Finally, another limitation of our work is that we treat genes independently, whereas it is more likely that the coordinated expression of a network of genes is more biologically relevant. Indeed, it was demonstrated that no single memory gene can predict the eventual persistence phenomenon in vemurafenib treated melanoma cells. Rather, groups of memory genes were required to accurately predict cell fate [16]. Constructing a principled approach to identifying memory ‘modules’, therefore, remains an exciting direction for future work.

## 5 Methods

### Model of gene expression dynamics

We assume each gene fluctuates about the steady state with a characteristic time scale of relaxation. We, therefore, model the gene expression dynamics of each gene as an Ornstein-Uhlenbeck process

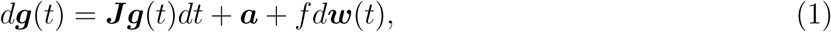

where ***g*** is a column vector for *n_g_* gene expression levels representing the concentrations of mRNA levels, ***J*** is a diagonal matrix whose elements denote the characteristic rates of relaxation for each gene (*µ_i_*), ***a*** is a column matrix with positive real numbers which determine the steady-state gene expression levels, ***w***(*t*) is the Wiener process and *f*, a positive real number, denotes the noise amplitude, which we take to be constant. We model the relaxation rates with a Gamma distribution *P* (*µ*; *φ, β*), parameterized by the shape *φ* and the rate *β*:

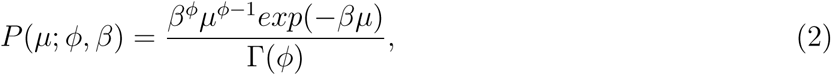

where Γ(*φ*) is the gamma function and *µ >* 0.

### Simulation of cellular lineages

We start with a single root cell with *n_g_* genes. The root cell undergoes a division process with a specified cell cycle time *τ_c_*. Each daughter cell perfectly inherits the gene expression of the mother since our expression measures the concentration. For each cell, we solve Eq. 1 to obtain the gene expression dynamics. We use the Euler-Maruyama method to numerically solve the stochastic differential equation.

In the simulation, ***a*** is a column matrix with dimensions (*n_g_,* 1) and all elements equal to 1. We use *f* = 0.1. Other values for *f* are shown in SI Fig. 3. For the relaxation rates modeled by a Gamma distribution *P* (*µ*; *φ, β*), we use *φ* = *β* = 2 if otherwise not mentioned. The initial state ***g*_0_** is a column matrix of dimension (*n_g_,* 1) with the entries sampled from a uniform random variable in the interval (0, 1). We equilibrate the gene expression of all the genes by waiting for time *t* = 1000 before starting the division process. If we have *b* divisions, the gene expression matrix (with dimensions (2*^b^, n_g_*)) is defined by the gene expression at the time step immediately prior to (*b* + 1)*^th^* division. The number of cells *n_c_*= 2*^b^*. To generate Fig. 2C-H, we have used *n_c_* = 2048, *n_g_* = 1024 and *τ_c_*= 0.5.

### Calculating the cell covariance matrix from simulations

From the gene expression matrix, we centralize each gene expression around its mean (*M_i_*) over all the cells, and then we normalize each gene expression value by its standard deviation across cells. This gives us the data matrix ***X****_S_*. Using ***X****_S_*, we calculate the cell covariance matrix ***C****_S_* = ***X****_S_*(***X****_S_*)*^T^ /n_g_*.

### Eigenspectrum of the true covariance matrix

The true cell covariance matrix is defined as the ensemble-averaged cell covariance matrix. Here, gene expressions are centralized about the ensemble mean and normalized by the ensemble standard deviation. If two cells are separated from their common ancestor by *m* cell divisions, the correlation between these two cells, as derived in the SI, is given by:

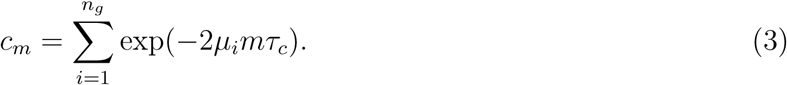

It is possible to derive an analytical expression for the eigenvalues of the true covariance matrix, generalizing the calculation in [31] (details in SI). If the division process has *b* branches (i.e, 2*^b^* cells), then the number of unique eigenvalues is (*b* + 1). These are:

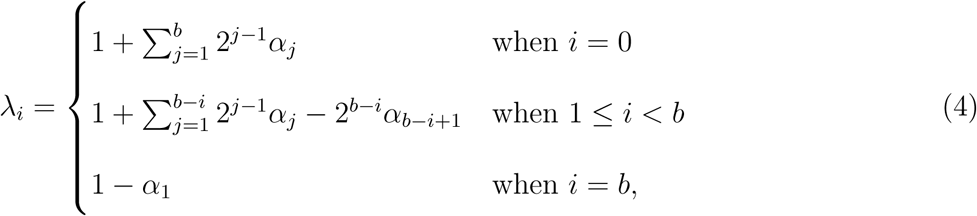

where *α_j_* = *c_j_/n_g_*. The eigenvalues are degenerate with the degeneracy equal to 1 (when *i* = 0) or 2*^i−^*^1^ (when 1 *≤ i ≤ b*). We exclude the largest eigenvalue which is not part of the power law and is given by the sum of the columns [31].

The mean half-life is defined as the number of generations it takes for the cell covariance to decay to half its initial value, and is therefore given by ln 2*/*(2*τ*), where *n_g_*is large and *τ* = *τ_c_/*(*φ/β*).

### Prediction of eigenspectrum of the single sample cell covariance matrix

The eigenspectrum of the sample covariance matrix can be predicted using the true covariance matrix [35, 31]. First, the distribution of eigenvalues of the true and sample covariance matrix can be related by doing a Stieltjes transform of the sample covariance matrix [35, 31]. It is possible to derive an algebraic equation for the Stieltjes transform which depends on the degenerate eigenvalues of the true covariance matrix. The Stieltjes transform can then be inverted to obtain the distribution of eigenvalues of the sample covariance matrix.

To find the analytical prediction for the ranked eigenvalues in Figure 2D, we use the mean order statistic of the distribution of the sample covariance matrix. Details of these calculations are given in the SI.

For large values of *τ*, the distribution of eigenvalues of the sample cell covariance matrix approaches the Marchenko-Pastur distribution, which is the predicted distribution for the sample covariance in the absence of any correlations. The Marchenko Pastur distribution is given by

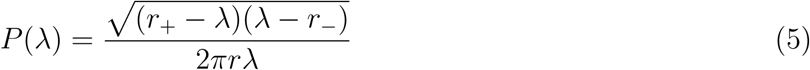

where 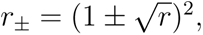 *r* = *n_c_*/*n_g_* and *r* < 1.

### Estimating the Power law exponent

We estimate the power law exponent by fitting a straight line to the top eigenvalues in the *λ*-vs-*R* curve on a log-log scale. We fit the straight line to a fraction of the eigenvalues. This fraction is determined by putting a threshold on the goodness of fit (by calculating *R*^2^-value). This threshold is set to 0.992 to generate Fig. 2 and Fig. 3, and 0.92 for Fig. 4.

### Shuffling

For every gene, we swap the expression state for the gene in a pair of cells that are randomly chosen. We repeat this 1.5 *× n_c_* times, where *n_c_* is the number of cells. The shuffled gene expression matrix is processed in the same way as the gene expression matrix to calculate the cell covariance matrix.

### Power-Seek algorithm

For the simulations, we remove the column corresponding to gene *g* from the gene expression matrix and calculate the updated data matrix 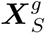 of order (*n_c_, n_g_ −* 1). We then calculate the updated cell covariance matrix, with the gene *g* removed 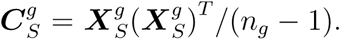. Therefore, we define a metric *δ_g_* for a specific gene *g* as 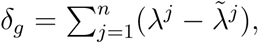, where *λ_j_* is *j^th^* rank eigenvalue of ***C****_S_* and 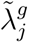 is *j^th^* rank eigenvalue of 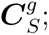; *n* is the number of eigenvalues of ***C****_S_* in the power law regime. Sorted values of *δ_g_* give us our predicted ordering of relaxation time scales with higher positive values of *δ_g_* corresponding to memory genes.

### Single cell RNA Data processing

We use scRNA data on the WM989 melanoma line from Harmange et al. [36] and on the human breast cancer tissue data [37]. We preprocess the data using Seurat V5 [50]. We filter cells based on the high percentage of mitochondrial and ribosomal genes and the number of unique genes in each cell.

In the WM989 cell line data set, of the two replicates available, we use one replicate to generate Fig. 4. The analysis for the other replicate is shown in Fig. SI 6. After following the preprocessing steps in Seurat, we remove genes that are not expressed in any cells in the data matrix. Reverse transcription and sequencing of a molecule can add variability to the measured raw UMI count in otherwise identical cells in scRNA-seq experiments. To minimize this variability, we calculate the total raw counts of each cell. Then, we divide the raw UMI counts for each gene in a cell with the total count of that cell. Sparsity can also affect the eigenvalue spectrum (Fig. SI 7). We, therefore, remove genes expressed in less than 200 cells. We also remove genes that have high expression in a small number of cells and low expression in all other cells using an outlier detection method (Fig. SI 5). Then, we select the top 5000 genes by comparing the coefficient of variations across cells and obtain the experimental raw count matrix of order (*n_c_, n_g_*), where *n_g_* = 5000. In case of WM989 cell line, *n_c_* = 7590 for replicate 1 and 4941 for replicate 2. For human breast tissue data, *n_c_* is 7967 and *n_g_* is 5000.

We then center the gene expressions about their mean and normalize each column by its standard deviation of gene expression over all cells to calculate an (experimental) data matrix ***X****_E_*. The experimental cell covariance matrix is defined as 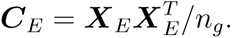. We use the Power-Seek algorithm described above to find *δ_g_*. To find the number of eigenvalues following the power law, we kept *R*^2^ = 0.92 for the WM989 cell line and 0.96 for Human breast cancer tissue data giving 50 eigenvalues in both cases.

### Overrepresentation analysis

We do the overrepresentation analysis on two datasets - a) top 200 Power-Seek rank genes from the WM989 cell line [36] and b) top 200 Power-Seek rank genes from human breast cancer tissue data [37], with a web-based software *WebGestalt* [38]. In each case, we use ‘cellular component’ and ‘molecular function’ as functional databases. We compare the representation of functional categories in memory genes with that in the top 5000 highly variable genes that we include in our data. If these genes are not annotated in the database, we ignore them.

For each dataset, we take the top 10 functional categories sorted by their p-value from low to high, calculated by *WebGestalt*. Then, we check the number of genes in a functional category in the reference and the number of genes in that functional category in the top 200 Power-Seek genes. We can calculate the probability of getting the set of genes in the top 200 Power-Seek gene list just by chance:

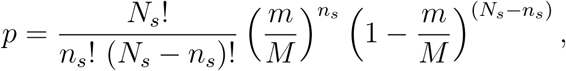

where *p* is a binomial probability, *N_s_*is the number of functionally annotated genes in the top 200 Power-Seek genes, *n_s_* denotes the number of genes in a specific functional category (say, F) among *N_s_* genes, *M* is the number of functionally annotated genes in the top 5000 high variable Power-Seek genes and *m* is the number of genes in F among *M* genes.

## Data and code availability

All original code has been deposited to GitHub at Cellmem-Power-Seek repo (click here). Github link: https://github.com/RajuLab/CellMem-Power-Seek

A corresponding Zenodo repository can be found under DOI: https://doi.org/10.5281/zenodo.20355272. Any additional information required to reanalyze the data reported in this paper is available from the lead contact(s) upon request.

## Acknowledgments

We acknowledge support from the Department of Atomic Energy, Government of India (under project RTI4006) and the Simons Foundation (287975). We thank Yuichi Wakamoto and Erik van Nimwegen for helpful comments.

